# CONSERVED INTERACTIONS IN CARDIAC SYNTHETIC THICK FILAMENTS DIFFERENTLY AFFECT MYOSIN SUPER-RELAXED STATE IN HEALTHY, DISEASE AND MAVACAMTEN-TREATED MODELS

**DOI:** 10.1101/2020.08.06.233213

**Authors:** Sampath K. Gollapudi, Suman Nag

## Abstract

A hallmark feature of myosin-II is that it can spontaneously self-assemble into bipolar synthetic thick filaments (STFs) in low ionic strength buffers, thereby serving as a reconstituted in vitro model for muscle thick filament. While these STFs have been extensively used for structural characterization, their use for functional studies has been very limited. In this report, we show that the ultra-low ATP-consuming super-relaxed (SRX) state of myosin is electrostatically more stable in STFs as compared with shorter myosin sub-fragments that lack the distal tail required for thick filament assembly. However, this electrostatic stability of the SRX state is weakened by phosphorylation of myosin light chains or the hypertrophic cardiomyopathy-causing myosin R403Q mutation. We also show that ADP binding to myosin depopulates the SRX population in STFs made of wild-type (WT) myosin, but not in S1, HMM, or STFs made of mutant R403Q myosin. Collectively, these findings emphasize that a critical network of inter- and intra-molecular interactions that underlie the SRX state of myosin are mostly preserved in STFs, establishing it as a native-like tool to interrogate myosin regulation. Next, using STFs, we show that a clinical-stage small molecule inhibitor, mavacamten, is more effective in promoting the myosin SRX state in STFs than in S1 or HMM and that it is equally potent in STFs made of atrial-WT, ventricular-WT, and mutant-R403Q myosin. Also, we found that mavacamten-bound heads are not permanently protected in the SRX state but can be recruited in response to physiological perturbations, thus providing new insights into its inhibitory mechanism.

## INTRODUCTION

Vertebrate striated muscle contraction is a result of cyclic interactions between myosin heads on the thick filaments and actin monomers on the thin filament. Emerging evidence in the last decade suggests that, in addition to Ca^2+^-mediated regulatory mechanisms within the thin filaments, the strength of muscle contraction may also be tuned by mechanisms intrinsic to myosin on the thick filaments (reviewed in (1–6)). Specifically, myosin heads in relaxed thick filaments are thought to exist in an equilibrium between two functional states (7–9): (1) the disordered relaxed state (DRX), in which myosin is free and ready to interact with actin and has an average ATP turnover time of <10 s; and (2) the super-relaxed (SRX) state, in which myosin is unavailable for interaction with actin and has a prolonged ATP turnover time of >100 s. Many studies have started showing that the availability of the myosin heads to form actin cross-bridges can be controlled by altering this DRX-SRX equilibrium, which in turn can regulate the strength of striated muscle contraction, all of which has been discussed in a recent report by Nag et al. ((7)).

The original discovery of the SRX state was based on single ATP turnover experiments in skinned rabbit fast and slow skeletal muscles (10), which was confirmed later by studies in skeletal and cardiac muscle systems from other species (8, 11–18). The structural basis for this biochemical SRX state is unclear due to a lack of high-resolution atomic-level structure of the myosin. At best, the functional SRX state of myosin is loosely correlated to the structural interacting-head motif (IHM) state of myosin in which the two myosin heads interact with one another while folding back onto their proximal tails (reviewed in (1–3)). In the muscle thick filaments, it has been hypothesized that, in addition to myosin interactions with titin and myosin-binding protein C (MyBPC), the folded-back myosin IHM state is stabilized by several intramolecular interactions among various sub-domains of myosin such as the regulatory light chain (RLC), heavy meromyosin (HMM), light meromyosin (LMM), subfragment-1 (S1) and subfragment-2 (S2), that give rise to S1-S1, S1-S2, RLC-RLC, S1-LMM and S2-LMM interactions (reviewed in (1–3)). If these interactions underlying the IHM state also give rise to the functional SRX state, then it is essential to build an experimental model that can capture most of these features. In previous studies, shorter myosin models such as full-length HMM, 25-hep HMM (HMM containing 25 heptad repeats of the S2), 2-hep HMM (HMM containing 2 heptad repeats of the S2), and S1 were shown to form the SRX state (13, 19). However, not only the SRX population was higher in full-length HMM and 25-hep HMM, but also the DRX-SRX equilibrium was more sensitive to electrostatic perturbations in 25-hep HMM than in either 2-hep HMM or S1 (13, 19). These observations suggested that the myosin SRX state is more stable in the presence of two S1 heads and the extended portion of the proximal tail. Importantly, myosin under physiological conditions does not exist in these short forms but present in an extensive filamentous form that is formed by the coiled-coil interactions of distal tails, highlighting an essential role for the extended tail in the myosin function. The present study focuses on employing reconstituted thick filaments, that mimics the structural environment close to the native thick filaments, for functional characterization of myosin in an in vitro setting.

There are two different ways to study myosin thick filaments. The first involves thick filaments extracted from the native muscle using gelsolin to remove the thin filaments (20, 21), and the second involves spontaneous self-assembly of myosin into bipolar thick filaments by lowering the ionic strength of the buffer containing full-length myosin ((22), reviewed in (23)). Of these two methods, the first one provides a native-like environment to study the myosin thick filament in the presence of MyBPC, titin, and other thick filament-associated proteins. The latter represents a simpler model that captures essential interactions pertinent to myosin alone while retaining the 14.3-nm myosin subunit periodicity and a 43-nm axial periodicity as in native filaments (24, 25). This model also provides a bottom-up approach to study the underlying biology of thick filaments and their function, starting from full-length myosin, offering advantages in different contexts. For example, this model permits the controlled addition of binding partners such as MyBPC to build a more complex system (26). Also, this system allows us to recombinantly study mutations in the rod domain of the myosin that causes several skeletal and cardiac myopathies (27, 28). Lastly, different studies have proposed co-operative myosin activation in thick filaments by several physiological perturbations such as ADP/ATP ratio (10, 15, 29), RLC phosphorylation (reviewed in (3)), actin and Ca^2+^ binding (30–32), which can be easily studied in a system containing assembled myosin filaments that preserves most of the myosin molecular interactions typically present in thick filaments. Despite this increasing need, the use of thick filaments for functional studies, especially in relevance to the low-energy consuming SRX state of myosin has been sparse.

In this study, using synthetic thick filaments (STFs) made from bovine and porcine cardiac full-length myosin, we tested the hypothesis that several intra-molecular interactions within the myosin molecule that are known to stabilize the IHM state of myosin may also underlie the SRX state. We provide evidence that the SRX population in STFs is electrostatically more stable as compared to that in shorter myosin subfragments, HMM, or S1. Multiple recent studies have demonstrated that a cardiac-specific myosin inhibitor (33, 34), mavacamten, which is currently in phase-III clinical trials for treating hypertrophic cardiomyopathy (HCM), reduces the cardiac muscle contractility by promoting myosin more into the SRX state (13, 19, 35). In support of this finding, using concentration-dependent responses in this study, we show that mavacamten promotes myosin in the SRX state with higher potency in STFs than in smaller myosin subfragments, HMM, and S1. However, we found that potency of mavacamten in promoting myosin into the SRX state is similar in STFs made of healthy atrial, healthy ventricular, or HCM-causing mutant (R403Q) myosin. We further demonstrate that the mavacamten-bound SRX heads in STFs can be reversibly recruited into the DRX state via the ADP-mediated cooperative activation of thick filaments, a mechanism that is not reproduced using HMM or S1.

## RESULTS

### Steady-state Measurements of Basal Myosin ATPase Activity in S1, HMM, and STF

To evaluate whether bovine cardiac (Bc) S1, HMM, and STF differ in their biochemical properties, we first measured the basal ATPase activity in these myosin systems. The resulting measurements did not show significant differences among the three myosin systems (Fig. 1A). The basal ATPase activities for BcS1, BcHMM, and BcSTF were 0.018±0.02 (*n*=8), 0.017±0.001 (*n*=6), and 0.016±0.001 (*n*=8) s^-1^, respectively. Similarly, the basal ATPase measured in other myosin models such as porcine cardiac (Pc) STFs made of WT and R403Q myosin in this study was also in the range of 0.010-0.025 s^-1^ (data not shown). Next, we measured basal ATPase activity in various myosin systems in response to increasing concentrations of mavacamten. A comparison of these responses did not show any obvious differences between systems in the potency of mavacamten on myosin ATPase inhibition (Figure 1B). Statistical analysis using one-way ANOVA confirmed that the concentrations of mavacamten required to attain half-maximal inhibition (IC_50_) in the basal ATPase activity were not significantly different between the three myosin groups. The IC_50_ values in BcS1, BcHMM, and BcSTF were 0.76±0.06 (*n*=8), 0.57±0.06 (*n*=6), and 0.82±0.06 μM (*n*=8), respectively, which are consistent with those reported for S1 and HMM in previous reports (19, 33, 34). These results suggest that BcSTF behaves the same way as BcHMM or BcS1 in the enzymatically coupled ATPase experiments (13, 19, 33, 34).

**Figure 1.**
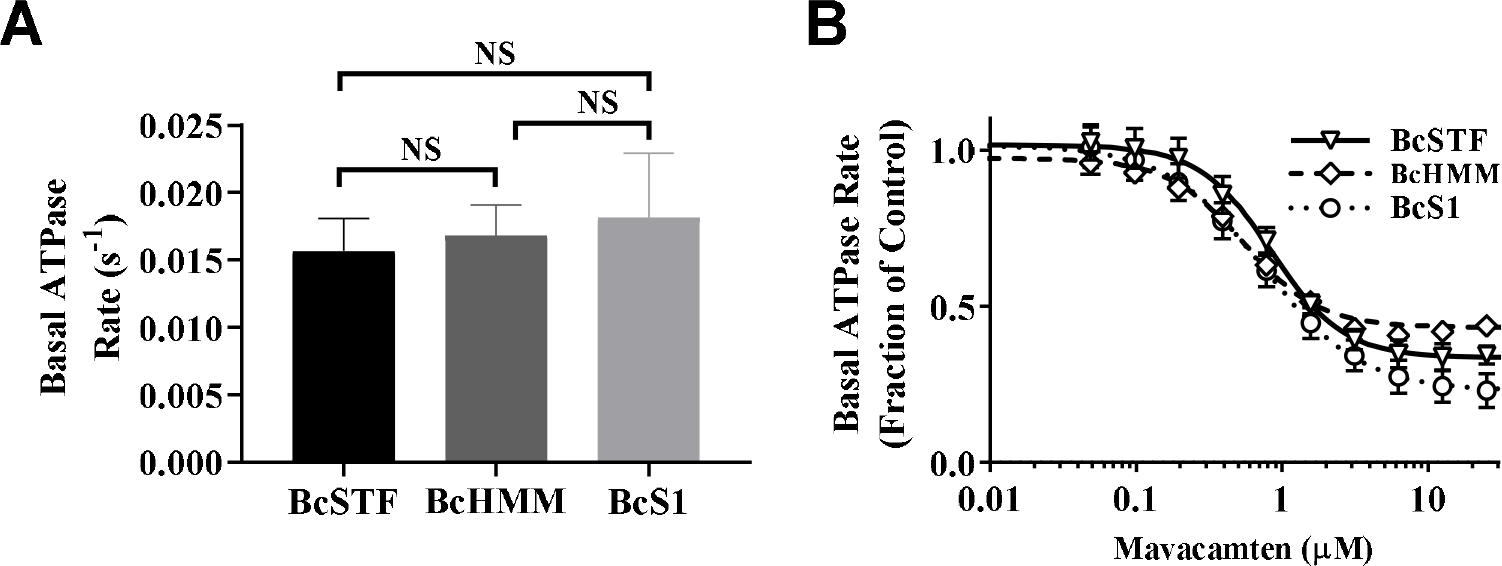
Basal ATPase of BcSTF, BcHMM, and BcS1 in the absence and presence of mavacamten. (A) Absolute basal ATPase rate (in *s*^-1^) of BcSTF, BcHMM, and BcS1 before treatment with mavacamten (NS, not significant). (B) Normalized basal ATPase profiles of BcSTF, BcHMM, and BcS1 following treatment with increasing concentrations of mavacamten. BcS1 refers to bovine myosin subfragment-1, BcHMM refers to bovine cardiac heavy meromyosin, and BcSTF refers to bovine cardiac synthetic thick filaments. The DMSO values were used to normalize the data in the respective untreated systems. The half-maximal change (IC_50_) of mavacamten estimated from these curves were 0.76±0.06 (*n*=8), 0.57±0.06 (*n*=6), and 0.82±0.06 μM (*n*=8) for BcS1, BcHMM, and BcSTF, respectively.

### Effect of Ionic Strength on Single Turnover Kinetics in S1, HMM, and STF

It is well established that the IHM state of myosin, which could also underlie the SRX state, involves a complex set of intra-molecular interactions that are highly electrostatic in nature (3, 36, 37). Therefore, in the next series of experiments, we perturbed these electrostatic interactions in BcS1, BcHMM, and porcine cardiac (Pc) STF by progressively increasing the ionic strength (KCl) of the buffer and examined the effect on parameters derived using single ATP turnover kinetic experiments. PcSTF was chosen in these experiments over BcSTF due to the ready availability of this reagent. However, it is unlikely that the use of PcSTF over BcSTF would have affected the conclusions drawn here, given the high sequence identity (~98%) between bovine and porcine β-cardiac myosin.

The fluorescence decay profile obtained during the chase phase in the single ATP turnover kinetic experiments characteristically depicted two phases, a fast phase followed by a slow phase (Fig. S1 in SI). Therefore, a bi-exponential function was fitted to estimate four different parameters—A_fast_, *k*f_ast_, A_slow_, and *k*_slow_—where A represents the % amplitude, and *k* represents the observed ATP turnover rate of each phase (10, 13, 19). Afast and *k*fast characterize myosin in the DRX state, while A_slow_ and *k*_slow_ characterizes it in the SRX state.

Analysis of data indicated that the response of PcSTF and BcHMM to increasing concentration of KCl was distinct from that of BcS1. For example, A_slow_, which represents the % amplitude of the myosin population in the SRX state, progressively decreased with increasing KCl concentration in both PcSTF and BcHMM but not in S1 (Fig. 2A). Importantly, PcSTF showed an even higher sensitivity of A_slow_ to ionic strength when compared to BcHMM. For instance, an increase in KCl concentration from 15 to 150 mM led to a net decrease in A_slow_ by 35±1% in PcSTF, as compared with only 20±5% in BcHMM. Besides, at a KCl concentration of 15 mM, A_slow_ was significantly higher by 14±5% in PcSTF than in BcHMM, suggesting that the tendency for myosin to form the SRX state at lower ionic strength is higher when myosin is assembled into the thick filament form. A similar dependency of myosin SRX population on ionic strength was also observed in rabbit cardiac myofibrils (Fig. S2 of Supporting Information), suggesting that the myosin in the SRX state is electrostatically better stabilized in STFs much in a similar manner as the native system. In contrast, myosin turnover rates in DRX (*k*_fast_) and SRX (*k*_slow_) states did not show any significant changes with ionic strength across all myosin types (Fig. 2B, 2C; Table S1 in SI). Collectively, these results suggest that the ionic strength alters the myosin DRX-SRX equilibrium without changing the intrinsic rates of myosin heads in these states.

**Figure 2.**
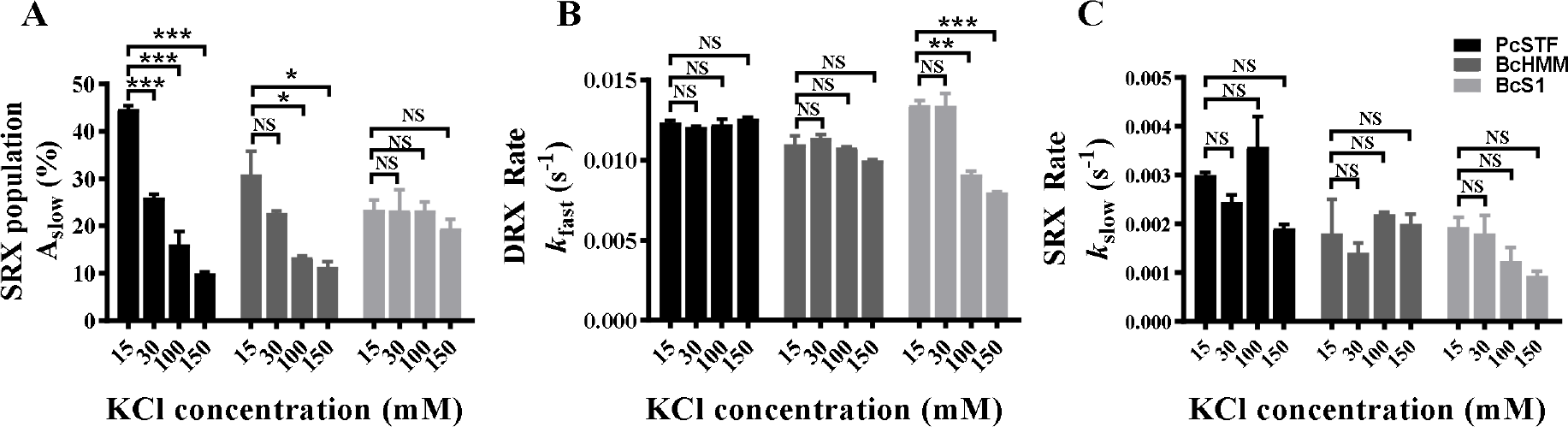
Effect of ionic strength (KCl) on the parameters derived from single ATP turnover experiments using BcS1, BcHMM, and PcSTF. Bar graphs showing the differences between PcSTF, BcHMM, and BcS1 at various ionic strengths for (A) A_slow_ (B) A_fast_ and (C) A_slow_. Parameters are reported on an absolute scale (amplitudes in % and rates in s^-1^). BcS1 refers to bovine myosin subfragment-1, BcHMM refers to bovine cardiac heavy meromyosin, and PcSTF refers to porcine cardiac synthetic thick filaments. Statistical differences were based on two-way ANOVA and subsequent post-hoc Tukey’s pair-wise comparisons (\**P*<0.01; * \**P*<0.01; \*\*\**P*<0.001; NS, not significant). Data are expressed as mean±SEM (*n*≥4 for each).

### Effect of Myosin Dephosphorylation on Single Turnover Kinetics in STF

Many studies have suggested that phosphorylation of the myosin regulatory light chain (RLC) controls myosin head conformation and hence can alter the myosin DRX-SRX equilibrium (3). In contrast, the role of the essential light chain (ELC) and the impact of its phosphorylation in this regard remains unclear. Here, we sought to examine the effect of a change in the RLC/ELC phosphorylation on the myosin SRX state. As described in the SI, we first ran a Pro-Q diamond gel to assess the phosphorylation status of RLC/ELC in a porcine cardiac full-length myosin sample, which confirmed that both RLC and ELC are phosphorylated to some extent in the basal state (Fig. S3 in SI). We treated the myosin sample with lambda phosphatase, which significantly reduced the phosphorylation levels of both RLC and ELC, as evidenced by weak phospho-stained bands in the lambda phosphatase-treated sample (Fig. S3 in SI). STFs made of lambda phosphatase-treated myosin showed a significant net increase in A_slow_ by 20±4% when compared to STFs made of untreated myosin (Fig. 3 A), suggesting that dephosphorylation of RLC/ELC shifts the myosin DRX-SRX equilibrium more towards the SRX state. This increase in A_slow_ is also associated with a 2-fold increase in both *k*_fast_ and *k*_slow_ (Fig. 3B, 3C), which are small enough increases to hold significant biological relevance.

**Figure 3.**
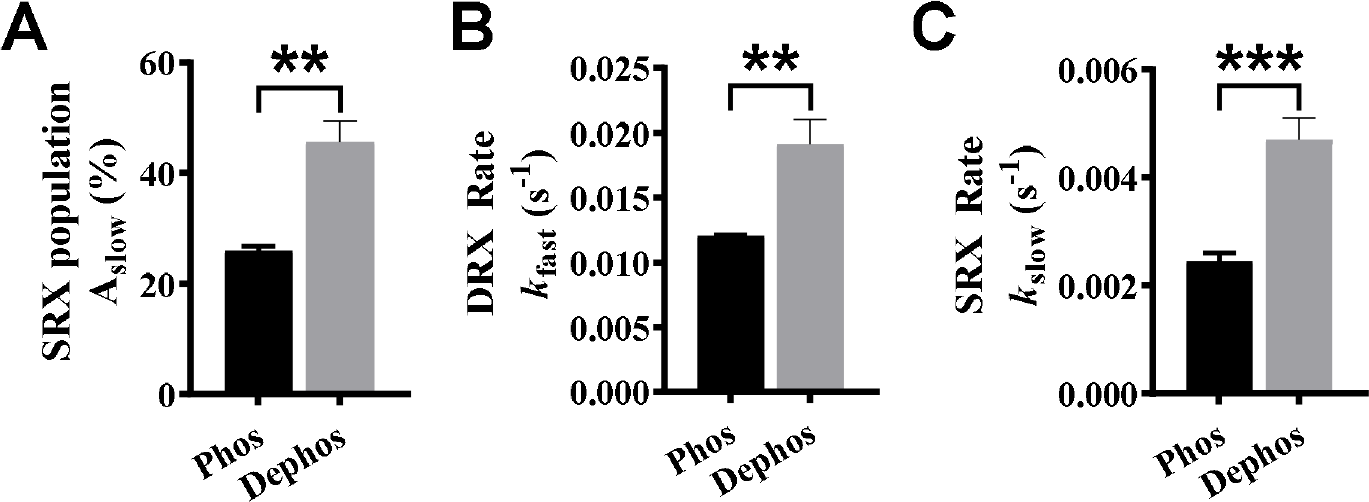
Effect of phosphorylation (Phos) and dephosphorylation (Dephos) on the parameters derived from single ATP turnover experiments using porcine cardiac synthetic thick filaments. Bar graphs showing the differences between untreated (with significant levels of phosphorylated ELC and RLC; Fig. S3 in SI) and dephosphorylated myosin samples for (A) A_slow_ (B) *k*_fast_ and (C) *k*_slow_. Parameters are reported on an absolute scale (amplitudes in % and rates in s^-1^). Statistical differences were based on two-tailed *t*-test (\**P*<0.01; \*\*\**P*<0.001). Data are expressed as mean±SEM (*n*≥4 for each).

### Effect of R403Q Myosin Mutation on Single Turnover Kinetics in STF

There is now substantial evidence from purified recombinant myosin to muscle fiber studies that the HCM-causing R403Q mutation in myosin shifts the equilibrium of myosin heads from the SRX state to the DRX state (13, 35, 38). We sought to test the same in STFs made of full-length myosin isolated from a porcine heart containing a heterozygous R403Q mutation. The STFs made are, therefore, likely to contain an equal mixture of the wild-type (WT) and R403Q myosin, similar to what may be present in a clinical situation. Changes in RLC phosphorylation are known to occur in disease hearts (39) and therefore, we first assessed the phosphorylation status of RLC and ELC in WT and R403Q myosin samples, as described in the SI. Our analysis showed that both the RLC and ELC are phosphorylated in either sample (see Fig. S3 in the SI). However, to avoid any contributions arising due to slight differences in the phosphorylation status, we completely dephosphorylated the WT and R403Q myosin samples using lambda phosphatase to normalize the phosphorylation status of RLC/ELC before conducting experiments (see Fig. S3 in the SI). We then measured single ATP turnover kinetics in STFs made from untreated and lambda phosphatase-treated WT and R403Q myosin at two different KCl concentrations (30 and 100 mM) and assessed the differences in A_slow_, *k*_fast_, and *k*_slow_. Post-hoc Tukey’s pair-wise comparisons from two-way ANOVA revealed that A_slow_ significantly decreased (*P*<0.001) from 46±4% in WT to 26±3% in R403Q group at 30 mM KCl, whereas it showed no significant change (*P*=0.83) between WT and R403Q groups at 100 mM KCl (Fig. 4A). In contrast, neither *k*_fast_ nor *k*_slow_ estimates were significantly different between groups at either KCl concentrations (Fig. 4B, 4C; Table S2 in SI). Our data in PcSTF verified that the R403Q mutation solely perturbs the SRX-DRX equilibrium to populate myosin heads more towards the DRX state, without affecting the intrinsic ATP turnover rate of myosin heads in the DRX and SRX states.

**Figure 4.**
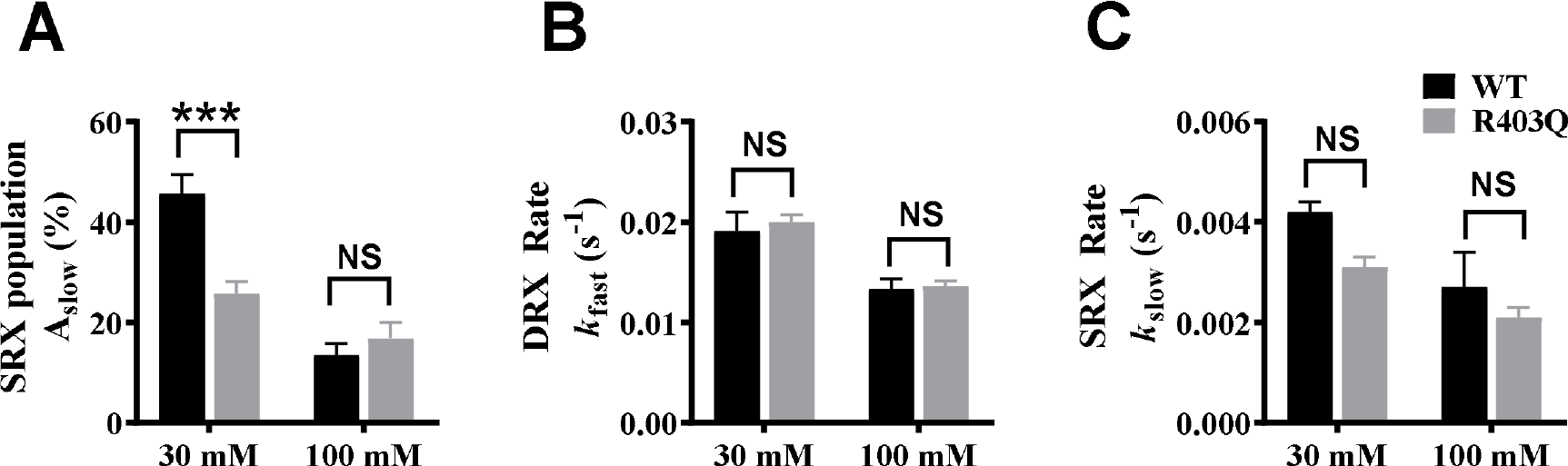
Effect of R403Q mutation in myosin on the parameters derived from single ATP turnover experiments using porcine cardiac synthetic thick filaments. Bar graphs showing the effect of R403Q mutation at 30 mM and 100 mM KCl concentrations for (A) A_slow_ (B) A_fast_ and (C) A_slow_. All parameters are reported on an absolute scale (amplitudes in % and rates in s^-1^). Statistical differences presented are based on two-way ANOVA and subsequent post-hoc Tukey’s pair-wise comparisons (\*\*\**P*<0.001; NS, not significant). Data are expressed as mean±SEM (*n*≥8 for each).

### Effect of ATP versus ADP Chase on Single Turnover Kinetics in Different Myosin Systems

Previous studies using single ATP turnover kinetics have shown that, under activating conditions, the slow phase (or the SRX phase) was diminished when cardiac and skeletal fibers incubated with mant-ATP were chased with ADP (10, 15, 29). To examine whether this phenomenon can be reproduced in simpler myosin systems, we performed ADP chase experiments in PcSTF-WT and PcSTF-R403Q systems. Our analysis showed that A_slow_ in PcSTF-WT significantly decreased from 46±4% when chased with ATP to 18±1% when chased with ADP (*P*<0.001; Fig. 5A) whereas this effect was not observed in PcSTF-R403Q (Fig. 5A). The decrease in A_slow_ observed in PcSTF-WT following ADP chase also coincided with a small but significant increase in *k*fast and a significant decrease in *k*slow (Fig 5B, 5C, and Table S2 in SI). However, as argued above, these changes in rates are too small enough to hold any biological relevance. Interestingly, neither BcS1 nor BcHMM showed any effect on A_slow_, *k*_fast_, and *k*_slow_ in response to ADP chase, when compared to ATP chase (Fig. S4 in SI). Collectively, these observations indicate that ADP-bound myosin heads also shift the DRX-SRX equilibrium more towards DRX state, but this effect requires a reserve in the SRX population as well as the presence of thick filament core, which are present in PcSTF-WT but not in BcHMM, BcS1 and PcSTF-R403Q (see Discussion).

**Figure 5.**
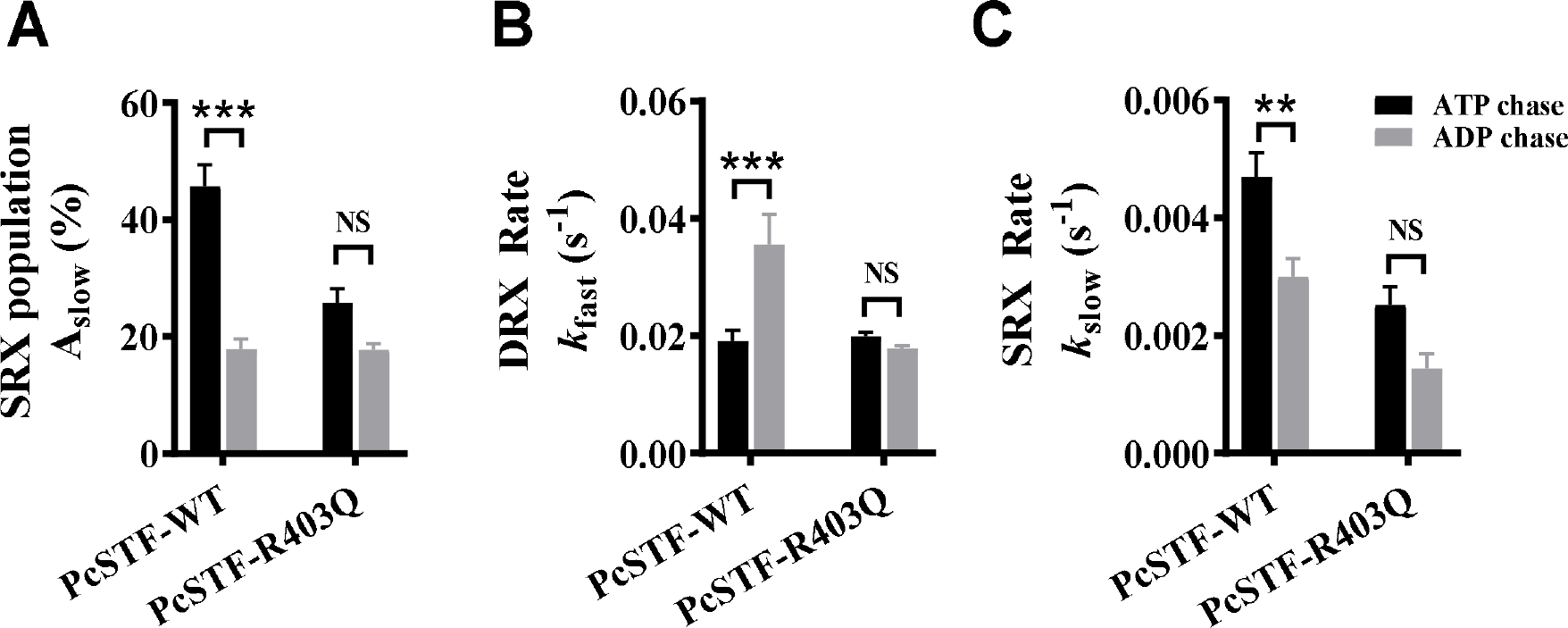
Effect of ATP chase versus ADP chase on the parameters derived from single ATP turnover experiments in STF models. Bar graphs showing the effect of ATP chase versus ADP chase in PcSTF-WT and PcSTF-R403Q for (A) A_slow_ (B) *k*_fast_ and (C) *k*_slow_. All parameters are reported on an absolute scale (amplitudes in % and rates in s^-1^). PcSTF-WT and PcSTF-R403Q refer to porcine cardiac synthetic thick filaments made of WT and mutant R403Q myosin. Statistical differences presented are based on two-way ANOVA and subsequent post-hoc Tukey’s pair-wise comparisons (\*\**P*<0.01; \*\*\**P*<0.001; NS, not significant). Data are expressed as mean±SEM (*n*≥6 for each).

### Effect of Mavacamten on Single Turnover Kinetics in S1, HMM, and STF

Previous studies have shown that mavacamten shifts the cardiac myosin DRX-SRX equilibrium more towards the SRX state (13, 19, 35). Here, we examined the concentrationdependent effect of mavacamten on the SRX state of STF and compared it with S1 and HMM. A comparison of resulting A_slow_ profiles showed a more prominent leftward shift in BcSTF than in BcHMM when compared to BcS1 (Figure 6A), suggesting that a lower concentration of mavacamten is required to attain the same increase in myosin SRX population in BcSTF than in BcHMM or BcS1. For example, at a concentration of 1.56 μM, mavacamten increased A_slow_ estimates in BcS1, BcHMM, and BcSTF to 36±1%, 70±10%, and 98±2%, confirming that mavacamten had a more significant effect in STFs over shorter myosin subfragments. This observation was supported by the EC_50_ values derived for A_slow_ in these systems. Post-hoc *t-*tests from one-way ANOVA showed that the EC_50_ of A_slow_ in BcSTF is significantly lower by 2-fold (*P*<0.05) when compared to BcHMM and by 3-3-fold (*P*<0.001) when compared to BcS1. The EC50 for Aslow in BcSTF, BcHMM, and BcS1 were 0.66±0.05, 1.17±0.13, and 2.16±0.19 μM, respectively. A similar trend was also noted in the ability of mavacamten in inhibiting *k*_fast_, the ATPase turnover rate of the myosin DRX heads (Figure 6B). The IC_50_ for *k*_fast_ in BcSTF, BcHMM, and BcS1 were 0.31±0.05, 0.54±0.08, and 0.67±0.09 μM, respectively. Collectively, these data indicate that the ability of mavacamten to promote more myosin into the SRX state is better realized in STFs than in shorter myosin models, S1 and HMM.

**Figure 6.**
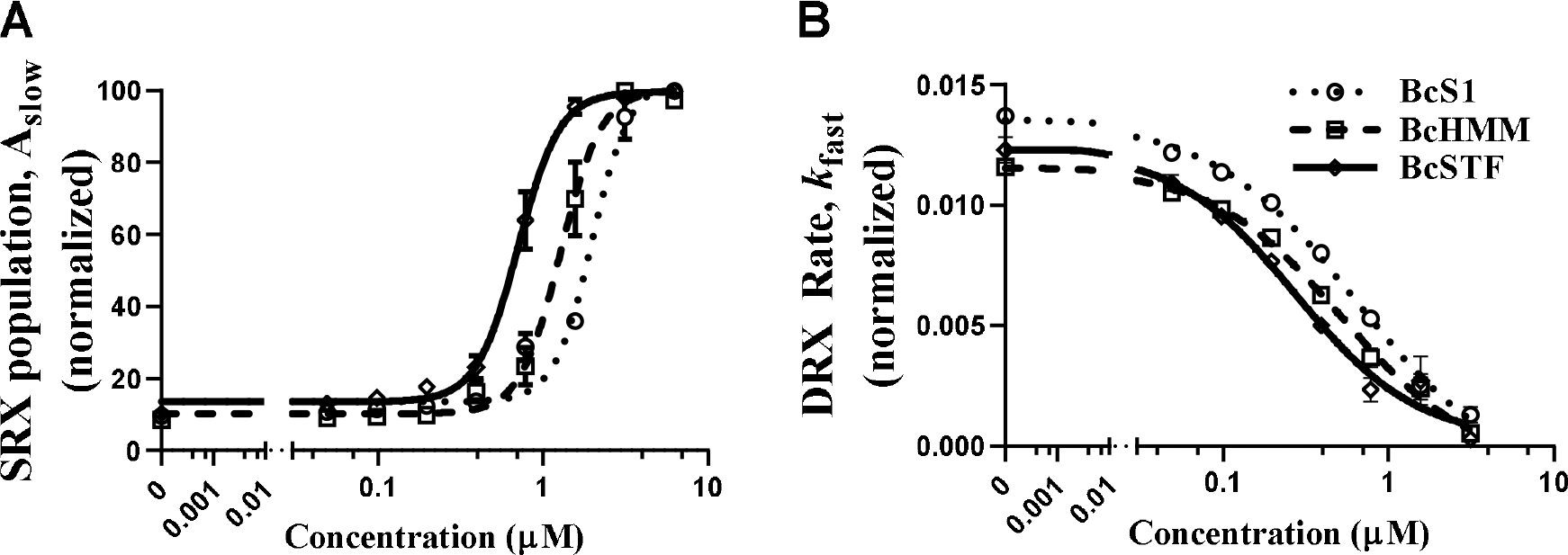
Effect of mavacamten on the parameters derived from single ATP turnover experiments in various myosin systems. Comparison of the responses of mavacamten between BcS1, BcHMM, and BcSTF for (A) A_slow_ and (B) A_fast_. BcS1 refers to bovine myosin subfragment-1, BcHMM refers to bovine cardiac heavy meromyosin, and BcSTF refers to bovine cardiac synthetic thick filaments. Data were expressed as mean±SEM (*n*=12 from three experiments). In panels A and B, concentrations of mavacamten required for a half-maximal increase in A_slow_ (EC_50_) for BcS1, BcHMM, and BcSTF were 2.16±0.19, 1.17±0.13, and 0.66±0.05 μM, respectively, while those required for the half-maximal decrease (IC_50_) in A_fast_ were 0.67±0.09, 0.54±0.08, and 0.31±0.05 μM, respectively.

### Effect of Mavacamten on Single Turnover Kinetics in STF Following ATP and ADP Chase

We next examined the effect of mavacamten on the SRX state in BcSTF in a concentration-dependent manner following ATP chase versus ADP chase and to tie its mechanism to the biological function. Figure 7A (open circles) shows that mavacamten populates myosin in the SRX state in a concentration-dependent manner, as evidenced by an increase in A_slow_. However, this ability to stabilize myosin heads in the SRX population was greatly diminished when chased with ADP (Fig. 7A; open triangles). The concentrations of mavacamten required for the half-maximal increase in myosin SRX population in the ADP chase experiments were significantly higher by 7-fold (*P*<0.001) when compared to those in ATP chase experiments. The EC_50_ of A_slow_ in the ATP chase and ADP chase experiments were 0.63±0.06 and >4.35±0.64 μM, respectively. Similarly, the ability of mavacamten to inhibit the DRX ATPase rate was significantly (*P*<0.05) reduced by 2.7-fold with ADP chase when compared to ATP chase. The DRX ATPase IC_50_ values for ATP chase and ADP chase experiments were 0.41±0.06 and 1.14±0.20 μM, respectively (Table S3 in SI). These results may hold significance in disease conditions such as HCM, where there could be a potential increase in the ADP concentration (40).

**Figure 7.**
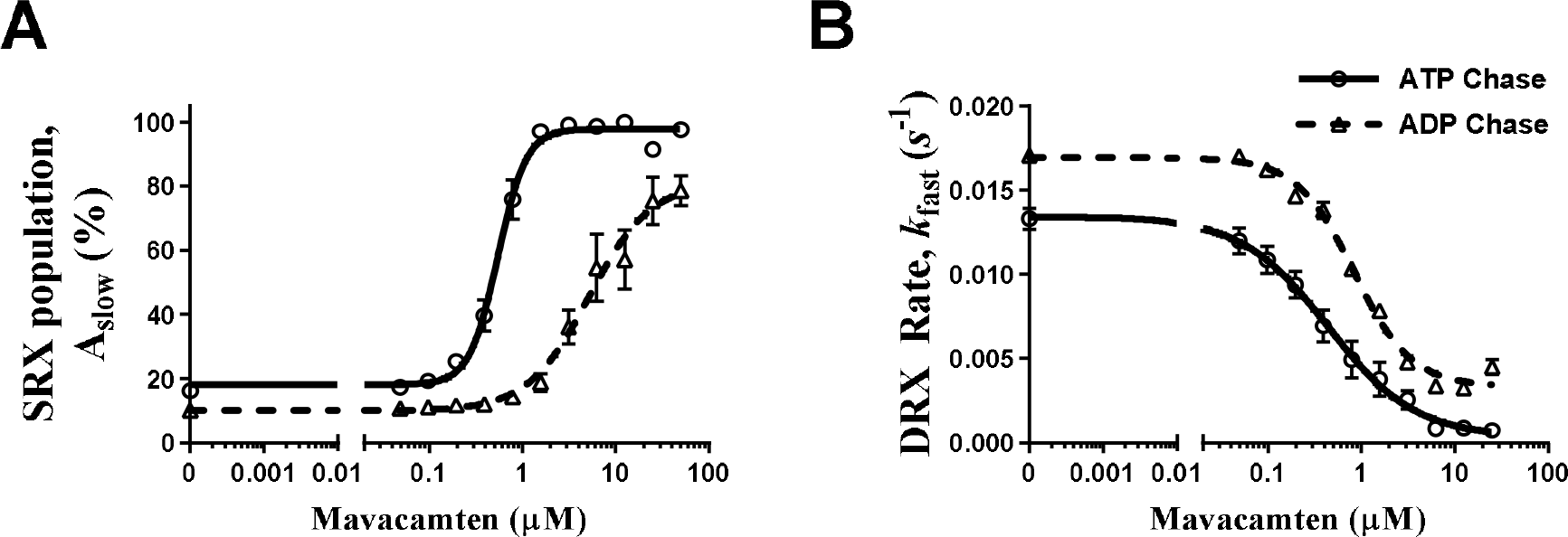
Effect of ATP vs. ADP chase in single ATP turnover experiments in the presence of mavacamten. Effect of ATP vs. ADP chase on (A) A_slow_ and (B) *k*_fast_ as a function of mavacamten. All experiments were performed using bovine cardiac synthetic thick filaments (BcSTFs). All parameters are reported on an absolute scale (amplitudes in % and rates in s^-1^). Data are expressed as mean±SEM (*n*≥4 from two experiments). The half-maximal change in A_slow_ for ATP chase and ADP chase experiments were 0.63±0.06 and >4.35±0.64 μM, while those for *k*_fast_ were 0.41±0.06 and 1.14±0.20 μM, respectively.

### Effect of Mavacamten in Healthy and Disease Models

Next, we compared the effect of mavacamten in healthy (PcSTF-WT) and disease (PcSTF-R403Q) in vitro model systems. As observed by an increase in Aslow, mavacamten increases myosin SRX population in a concentration-dependent manner with almost equal potency in either system (Fig. 8A). The concentrations required for a half-maximal increase in myosin SRX population were not statistically significant in WT and R403Q systems, and these were 1.76±0.22 and 2.00±0.28 μM, respectively. The increase in A_slow_ was associated with a concomitant decrease in *k*f_ast_ with no significant differences between groups. The IC_50_ estimates of *k*f_ast_ for WT and R403Q were 0.29±0.02 and 0.30±0.02 μM, respectively (Figure 8B and Table S3 in SI), which are supported by data from the basal ATPase assay (data not shown). These observations suggest that mavacamten inhibits myosin activity by effectively populating myosin in the SRX state even in a disease model.

**Figure 8.**
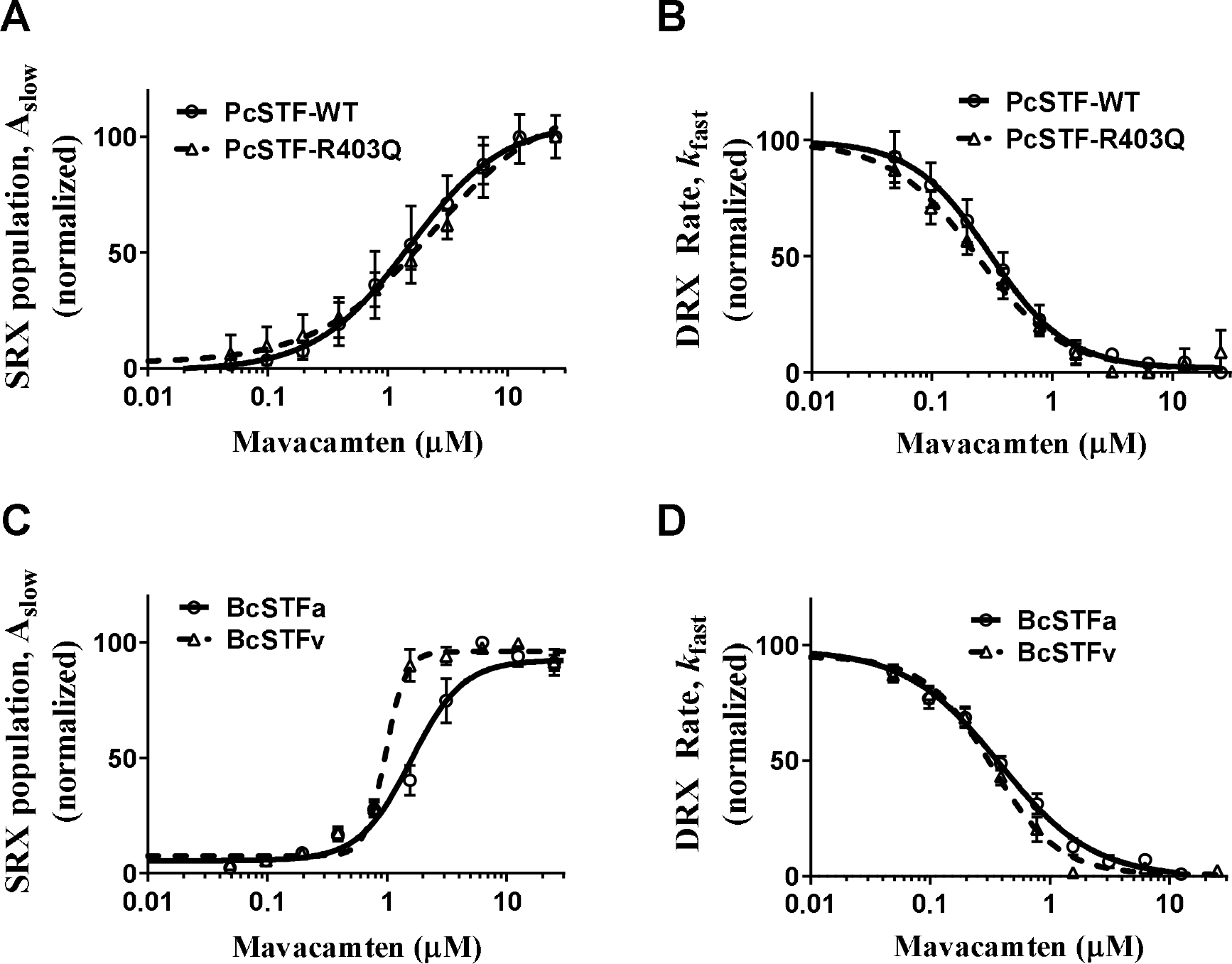
Effects of mavacamten in healthy and disease models. Comparison of the effect of mavacamten between PcSTF-WT and PcSTF-R403Q for (A) A_slow_ and (B) A_fast_. Comparison of the effect of mavacamten between BcSTFs made from α-cardiac and β-cardiac myosin for (C) A_slow_ and (D) A_fast_. Nomenclature is as follows: PcSTF and BcSTF refer to porcine and bovine cardiac synthetic thick filaments, and BcSTFa and BcSTFv bovine cardiac synthetic thick filaments made from left atrial and left ventricular full-length myosin, respectively. Data were expressed as mean±SEM (*n*≥4 from two experiments). In (A) and (B), the half-maximal change in A_slow_ for WT and R403Q PcSTF were 1.76±0.22 and 2.00±0.28, while those for A_fast_ were 0.29±0.02 and 0.30±0.02 μM, respectively. Similarly, in (C) and (D), the half-maximal change in A_slow_ for α-cardiac and β-cardiac BcSTF were 1.83±0.25 and 1.03 ±0.11 μM, while those for ft were 0.61±0.08 and 0.35±0.06 μM, respectively.

Recent studies have shown that in cellular models of MYH6 expressing the faster α-cardiac myosin, mavacamten inhibits myosin activity and populates the SRX state, similar to its effect in models expressing the slower β-cardiac myosin (35). However, there are no biochemical studies comparing the potency of mavacamten in these two systems side-by-side. To address this, we first purified myosin from the left atrial and ventricular chambers of the bovine heart, and using a native PAGE gel, we confirmed that they predominantly express faster α-cardiac and slower β-cardiac myosin isoforms, respectively (Fig. S5 in SI). We then made BcSTF using these myosin variants and conducted single turnover experiments in the presence of increasing concentrations of mavacamten. A comparison of concentration-dependent profiles of A_slow_ (Figure 8C) and *k*f_ast_ (Figure 8D) suggests a lower efficacy of mavacamten in both increasing A_slow_ and decreasing *k*_fast_ in BcSTFa (α-cardiac) than in BcSTFv (β-cardiac) systems. In support of this observation, our analysis confirmed that both the EC_50_ of A_slow_ and the IC_50_ of *k*f_ast_ in BcSTFa were significantly (*P*<0.05) higher by 1.8-fold in BcSTFa when compared to those in BcSTFv. The IC_50_ values of mavacamten for A_slow_ in BcSTFa and BcSTFv were 1.83±0.25 and 1.03 ±0.11 μM, while those for *k*f_ast_ were 0.61±0.08 and 0.35±0.06 μM, respectively (Figures 8C-D and Table S3 in SI). These data are consistent with the observed inhibition in the basal ATPase assay among different myosin systems following mavacamten treatment (data not shown). Taken together, these data suggest that mavacamten is slightly more effective in enhancing the SRX state in ventricular than in atrial myosin systems. Whether such small differences hold any relevance to clinical applications of mavacamten remains to be understood.

## DISCUSSION

Muscle myosin-II is long known to spontaneously form bipolar synthetic thick filaments (STFs) in buffers of low ionic strength ((22), reviewed in (23)) With some caveats, such reconstituted thick filaments have been shown to structurally mimic essential features similar to the assembled myosin in native thick filaments (23). However, other than myosin assembly and disassembly kinetics, these STFs have found minimal use for functional studies of myosin ((41–46) and reviewed in (23)). In this study, using STFs as our primary experimental model, we tested the hypothesis that the myosin folded-back interacting head motif (IHM) state, which is held by several inter- and intramolecular interactions between different sub-domains of myosin stabilizes it in an energy-sparing myosin super-relaxed state (SRX) population. Our findings demonstrate that STFs reproduce critical features of the myosin SRX state, many of which are missed in the truncated myosin systems such as S1 and HMM.

Basal myosin ATPase assays demonstrated that the ATPase activity was similar among bovine cardiac STFs, HMM, and S1. The efficacy of mavacamten in inhibiting basal myosin ATPase was also not different across these systems (Fig. 1). These results suggest that the overall ATPase activity of myosin was not altered in BcSTF despite holding a more complex structure as compared to BcHMM and BcS1. However, it is worth noting that the ATP turnover rate measured in the ATPase assay represents the ensemble-averaged rate of myosin heads in the DRX and SRX states but does not allow quantification of myosin population in these two states. Such quantification of myosin distribution in the SRX and DRX states requires single ATP turnover experiments. The biochemical SRX state, with an ultra-low ATP turnover rate, has been hypothesized to be related to the IHM state (9, 11, 47–49). In such a structural state, the two myosin S1 heads interact with one another and asymmetrically fold back onto their respective proximal S2 tails, giving rise to several intra-molecular interactions among different subdomains of myosin such as head-head (S1-S1), head-tail (S1-S2), RLC-RLC, S1-LMM, and S2-LMM (3).

Given that all such possible interactions are present in STFs, a few of them in HMM, and none in S1, we sought to validate the role of these electrostatic interactions in stabilizing the SRX state of myosin. Consistent with this notion, the myosin SRX population in STFs and HMM, but not in S1, showed a graded decrease in response to increasing ionic strength (Fig. 2). Our findings related to HMM and S1 are highly consistent with two previous studies (13, 19). More importantly, the magnitude of reduction in the myosin SRX population is higher in STFs than in HMM for an equivalent increase in ionic strength. Interestingly, much of this difference can be attributed to a higher myosin SRX population in STFs at the lowest ionic strength (15 mM KCl), when compared to HMM. Structurally, one crucial difference between myosin in STFs versus HMM is that the long distal tails of the full-length myosin in STFs can self-associate to form the thick filament backbone at low ionic strengths through intermolecular interactions such as S1-LMM, S2-LMM, LMM-LMM, etc. Therefore, our observations suggest that the thick filament backbone is essential to promote all inter- and intra-molecular interactions that stabilize myosin in the SRX state. This was also evident in the report by Anderson et al. (13), where the population of the SRX in the muscle fibers was higher than in purified proteins despite the high ionic strength used in these experiments. Similarly, consistent with other studies (10, 12, 31, 50), we show that RLC phosphorylation destabilizes the myosin SRX population in STFs (Fig. 3), further suggesting that electrostatic interactions play a crucial role in holding myosin population in the SRX and IHM states.

In addition to demonstrating the role of complex molecular interactions in the myosin SRX state in STFs, we also show that STFs made of full-length myosin purified from the left ventricular muscle of heterozygous R403Q HCM pigs have less myosin in the SRX state when compared to those made of WT (Fig. 4). This is consistent with the behavior observed in both purified myosin and muscle fibers containing the myosin R403Q mutation (13, 38). Interestingly, unlike WT, the SRX population in STFs made from R403Q heterozygous pigs did not show any ionic strength dependence from a statistical standpoint. It has been suggested in the literature that the R403Q mutation in myosin lies in the putative interface between the two heads, and hence it disrupts the SRX state by weakening the S1-S1 interaction, but not by altering S1-S2 interaction (36). Our results indicate that even at low ionic strength, R403Q weakens such S1-S1 interactions to facilitate myosin recruitment from the SRX to the DRX state, leaving behind a lower myosin SRX reserve that can be recruited by any mechanism. Consequently, STFs made of R403Q myosin elicited a small insignificant decrease (from 26±2% to 17±3%; Fig. 4A) in the SRX population in response to an increase in ionic strength. On the contrary, the initial SRX reserve in STFs made of WT myosin was higher, and therefore, the resulting decrease in the myosin SRX population to an increase in ionic strength was also significantly higher in this system (from 46±4% to 14±2%; Fig. 4A).

Single turnover experiments performed with slow skeletal, fast skeletal, cardiac, and tarantula muscle fibers have shown disruption of the myosin SRX state in the presence of ADP (10, 15, 29). At least with the skeletal muscle fibers, this observation has also been qualitatively recapitulated when studied at very long sarcomere lengths where there is no overlap between actin and myosin, suggesting that the binding of ADP to some myosin heads co-operatively disrupts the SRX property of the neighboring myosin heads through a pathway that resides within thick filaments. Very interestingly, such co-operative disordering (or activation) was suggested to be less prominent in cardiac than in fast skeletal muscle because ADP-bound myosin heads fully depopulate the myosin population from the SRX to the DRX state in fast skeletal but not in cardiac muscle (10, 11). Consistent with this hypothesis, STFs made of cardiac full-length myosin, but not S1, showed a significant decrease in the myosin SRX population but did not fully depopulate this state when chased with ADP versus ATP. This observation further bolstered the notion that ADP-bound myosin heads may recruit some myosin heads from the SRX to the DRX state via co-operative activation of the thick filament (Fig. 5). Such cooperative activation can only be achieved via a relay system that involves a communication between the proximal tails of the myosin to the distal tails that form the thick filament backbone, which is not present in HMM or S1. This may explain why the ADP chase did not affect the parameters of single ATP turnover experiments in either HMM or S1 (Fig. S4 in SI). On the contrary, STFs made of R403Q myosin did not show a significant change in the myosin SRX population when chased with ADP. As argued above, the ability of any perturbation to recruit myosin is a function of the initial myosin population in the SRX state. This fraction being lower in STFs made of R403Q myosin, the resulting decrease in the SRX population upon ADP chase was lower (from 26±2% to 18±1%; Fig. 5A) when compared to STFs made of WT myosin (from 46±4% to 18±2%; Fig. 5A).

It has been hypothesized that myosin heads in the SRX state act as a reserve and may be pulled into action in response to increased physiological demand (7–9). Along this line, it has been hypothesized and shown in many cases that several HCM-causing mutations in either myosin or MyBPC disrupt the SRX state and pull more myosin into play, resulting in clinical hypercontractility (7, 13, 18, 38, 51–54). To negate this effect, MyoKardia Inc. developed a small molecule cardiac myosin inhibitor, mavacamten (33, 34), which at a single high concentration, has been previously shown to shift the myosin DRX-SRX equilibrium towards the SRX state, thereby increasing the myosin SRX reserve (13, 19, 35, 51). By studying the concentrationdependent effect of mavacamten, we show here that it brings about inhibition in the basal ATPase activity (Fig. 1) by primarily increasing the myosin SRX population while concomitantly decreasing the observed ATP turnover rate of the DRX heads in all myosin systems evaluated in this study including BcS1, BcHMM, and BcSTF (Fig. 6). Our finding that mavacamten promotes a more significant myosin fraction from the DRX to the SRX state in HMM than in S1 is consistent with recent reports (13, 19). Interestingly, this ability of mavacamten in promoting myosin into the SRX state is further enhanced in BcSTF, reiterating the notion that all the key structural interactions that underlie the myosin SRX state are better preserved in STFs than in HMM or S1. However, our result that mavacamten can promote all the myosin from the DRX to the SRX state in S1 and HMM differ from previous studies ((19),(13)), but is consistent with the fact that mavacamten completely inhibits the basal ATPase activity in these systems (Fig. 1A).

To further demonstrate that mavacamten does not lock myosin heads permanently in the SRX state and that these heads can be recruited into the DRX state when required, we also performed ADP chase experiments (Fig. 7). Consistent with the activation mechanism detailed above for the ADP-bound myosin heads, mavacamten was less effective in increasing the occupancy of myosin population in the SRX state when chased with ADP at all concentrations (Fig. 7). This is substantiated by the 7-fold increase in the EC_50_ of mavacamten for myosin SRX population and 2.7-fold increase in the IC_50_ for the DRX ATPase rate in ADP chase experiments relative to those in ATP chase experiments. An alternate explanation could be that mavacamten may bind weakly to myosin in the myosin-ADP state, and consequently, this may explain a lower SRX population independent of mavacamten concentration. Collectively, these observations made in STFs not only validate the inhibitory mechanism of mavacamten but also confirm that the mavacamten-bound heads can be reversibly recruited into the DRX state when needed. Such ADP-triggered activation of the muscle and the reduced SRX stabilization by mavacamten may be relevant in disease conditions such as HCM or effects of inotropes that could potentially increase the [ADP]/[ATP] ratio (40).

For therapies involving the use of small-molecule drugs such as mavacamten, it is of considerable interest to evaluate the efficacy of these agents in a disease setting. Earlier, we have shown that the HCM-causing R403Q mutation induces a hypercontractile function by shifting the myosin DRX-SRX equilibrium more towards the DRX state, presenting an excellent HCM disease model in which mavacamten can be tested as a proof of concept. When tested in this disease model, we found that mavacamten is equally effective in populating the myosin SRX state in STFs made of WT and R403Q myosin (Fig. 8), consistent with a previous study showing the same in porcine ventricular fibers containing R403Q mutation at a single high concentration of mavacamten (13).

Clinical studies have shown that left ventricular diastolic dysfunction, a consistent characteristic feature of HCM (55), sets elevated left ventricular pressures against which left atrium has to fill, resulting in pathological hypertrophy and dysfunction of the left atrium (55). In clinical studies of mavacamten, it is observed that it reduces left atrium size, most likely by reducing left ventricular filling pressure and diastolic dysfunction (56). Given that mavacamten is not selective for the α-cardiac and β-cardiac myosin, theoretically, we would expect that mavacamten would have the same inhibitory ‘relaxing’ effect on the left atrium similar to that of the left ventricle. This ‘relaxing’ effect, which should enlarge rather than compact the left atrium, is compensated by the reduction in back-pressure from the left ventricle, leading to a more compact left atrium. Keeping up to this expectation, we observed that mavacamten equally populates the SRX state of the α-cardiac myosin purified from the left atrium as compared to those of the β-cardiac myosin purified from the left ventricle, with similar efficacies. This suggests that in a clinical setting, mavacamten may have a direct benefit to both atrial and the ventricular function.

In summary, we demonstrate here that reconstituted bipolar thick filaments formed from full-length myosin capture essential inter- and intra-molecular interactions between various subdomains of myosin, which are essential to stabilize the myosin SRX state. The interplay of these complex molecular interactions is necessary while understanding the effects of physiological and pathological perturbations in the cardiac sarcomere. The use of such reconstituted bipolar thick filaments has become more valuable for studies that focus on the co-operative mechanisms within thick filaments, a topic that has been mostly unexplored but gaining momentum in the last decade or so. For example, many biologically relevant perturbations such as Ca^2+^, ADP, disease-causing thick filament protein mutations, and possibly small molecules, as we show in this study can modulate the co-operative mechanisms within thick filaments, which can be better investigated using an experimental model that is close to the physiological setting. Therefore, the reconstituted bipolar thick filament model put forth here is not only expected to provide new avenues to advance our understanding of myosin structure-function relationships in a variety of contexts but may also aid us in the discovery of better therapeutic agents for disease prevention.

## MATERIALS AND METHODS

### Cardiac Protein Preparations

Bovine cardiac actin was purified following previously established method (57) with some modifications. Actin was stored at −80°C as G-actin and polymerized fresh for each day of experiments by adding 50 mM KCl and 2 mM MgCl_2_ to the actin-containing buffer. β-cardiac full-length myosin from bovine and porcine left ventricles and α-cardiac full-length myosin from bovine left atrium were isolated following established methods described elsewhere (58). Following this, proteins were dialyzed in a buffer containing 10 mM Pipes (pH 6.8), 300 mM KCl, 0.5 mM MgCl2, 0.5 mM EGTA, 1 mM NaHCO3, and 1 mM DTT and stored at −80°C. Using bovine cardiac full-length myosin as the starting material, heavy meromyosin (HMM) and sub-fragment S1 were prepared according to methods described elsewhere (58). Rabbit cardiac myofibrils, prepared using previously established methods (59), were dialyzed into PM12 buffer (12 mM PIPES, 2 mM MgCl_2_, pH 6.8) containing 10% sucrose for storage at −80°C.

### Reconstitution of Myosin Thick Filaments

Full-length myosin remains in a soluble form in a buffer of high ionic strength (300 mM), but many previous studies had shown that it could spontaneously self-assemble into bipolar thick filaments when the ionic strength of the buffer was reduced to 150 mM or below (22, 23). This method was used to construct thick filaments with some modifications in this study. Briefly, the ionic strength of the myosin sample, as well as the myosin concentration, were first adjusted to the desired value by diluting it with a buffer containing: 20 mM Tris-HCl (pH 7.4), 0 mM KCl, 1 mM EGTA, 3 mM MgCl_2_, and 1 mM DTT. After dilution, the myosin sample was incubated for two hours on ice to allow the formation of thick filaments before using it for experiments. In this study, such reconstituted myosin filaments will be referred to as synthetic thick filaments (STFs). Alternatively STFs were also made by slowly dialyzing full-length myosin into a low ionic (150 mM or below) strength buffer. Quantitatively, the measurements reported here were not different between STFs made by these two methods. Four different STF models were generated for this purpose, which includes STFs made of porcine WT ventricular myosin (PcSTF or PcSTF-WT), STFs made of porcine R403Q mutant ventricular myosin (PcSTF-R403Q), STFs made of bovine WT ventricular myosin (BcSTF or BcSTFv), and STFs made of bovine WT atrial myosin (BcSTFa).

### Steady-state ATPase Measurements

Measurements of basal myosin ATPase activity in BcSTF, BcHMM, and BcS1 systems as a function of increasing mavacamten concentrations were performed at 23°C on a plate-based reader (SpectraMax 96-well) using an enzymatically coupled assay (34). The composition of the buffer used for these experiments was 12 mM Pipes (pH 6.8), 2 mM MgCl2, 10 mM KCl, and 1 mM DTT. A concentrated stock (20 mM) of mavacamten, synthesized by MyoKardia, Inc., was first prepared using dimethyl sulfoxide (DMSO), and this was used to attain the desired concentration ranging from 0 to 50 μM in the final buffer samples. A final DMSO concentration of 2% was achieved in all samples. Data recorded by the instrument compatible software package, SoftMax Pro, was exported to GraphPad Prism and analyzed. A final myosin concentration of 1 μM was attained irrespective of the myosin model used.

### Single mant-ATP turnover Kinetics

Single ATP turnover kinetics with a fluorescent 2’/3’-O-(N-Methylanthraniloyl) (mant)-ATP were measured using a two-step mixing protocol using methods described previously (10, 13, 19). The myosin kinetics of this fluorescent mant-ATP is very similar to that of the non-fluorescent ATP (60). In the first step, myosin was combined with mant-ATP in a 1:1 ratio, and the reaction was aged for 60 s to allow binding and hydrolysis of mant-ATP to inorganic phosphate (Pi) and mant-ADP. In the second step, mant-nucleotides were chased with non-fluorescent ATP by adding an excess of non-fluorescent ATP to the mixture, and the resulting fluorescence decay due to mant-nucleotide dissociation from myosin was monitored over time. The composition of the final buffer was as follows: 20 mM Tris-HCl (pH 7.4), 30 mM KCl, 1 mM EGTA, 3 mM MgCl_2_, and 1 mM DTT. The concentrations of myosin, mant-ATP, and non-fluorescent ATP attained in the final mixture were 125 nM, 125 nM, and 4 mM, respectively. The excitation wavelength for monitoring the mant-nucleotides was 365 nm, and the emission wavelength was at 450 nm. Data were acquired at a frequency of 60 Hz. All experiments were performed at 25°C.

Consistent with similar studies in the past (10, 13, 19), the fluorescence decay profile obtained during the chase phase characteristically depicts two phases, a fast phase followed by a slow phase (Fig. S1 in SI). Therefore, a bi-exponential function was fitted to each trace to estimate four parameters corresponding to fast and slow phases — A_fast_, *k*_fast_, A_slow_, and *k*_slow_ — where A represents the % amplitude and *k* represent the observed ATP turnover rate of each phase (Fig. S1 in SI). In this study, the fast and the slow phases correspond to the myosin activity in the DRX and SRX states, respectively.

### Data Analysis

For each assay, the experiment was repeated at least twice with a minimum of two replicates per experiment. Sample number *n* refers to the total number of measurements made (number of experiments*number of replicates), which were averaged and presented as mean ± SEM. Dose-response profiles obtained using mavacamten were used to estimate the concentration (in μM) required to attain half-maximal change (IC_50_) in a given parameter. In experiments involving only one factor, we used a two-tailed student’s *t*-test. For experiments involving two factors, we used a two-way ANOVA to analyze the effect of both factors. Subsequent post-hoc *t*-tests using Tukey’s pair-wise comparisons were used to differentiate the changes in parameters among groups statistically. Significance was set at *P* < 0.05.

## DATA AVAILABILITY

All data presented in this manuscript is either included in the main article or the Supporting Information.

## ACKNOWLEDGMENTS

The authors would like to thank the protein production team and the medicinal chemistry team of MyoKardia Inc. for the production of biological reagents and mavacamten, respectively. The authors thank Ming Yu of Myokardia Inc. for his help in developing the SRX assay. The authors also thank Roger Cooke, Professor Emeritus at UCSF; Robert McDowell, CSO of MyoKardia Inc.; Fen Gan, Robert Anderson, and Uwe Klein, Biology leaders at MyoKardia Inc. for their critical review and suggestions in improving the manuscript.

