## Supplemental information for "CONSERVED INTERACTIONS IN CARDIAC SYNTHETIC THICK FILAMENTS DIFFERENTLY AFFECT MYOSIN SUPER-RELAXED STATE IN HEALTHY, DISEASE AND MAVACAMTEN-TREATED MODELS"

Running title: Super-relaxed state in reconstituted myosin thick filaments

Keywords: Super-Relaxed State (SRX), Synthetic thick filaments (STF), mavacamten, R403Q, Interacting Head Motif (IHM)

**MATERIALS AND METHODS**

#### **Determination of Phosphorylation Status of Thick Filament Proteins**

The concentrations of wild-type (WT) and mutant (R403Q) full-length myosin samples were first adjusted to 2 mg/ml in a buffer containing Pipes (pH 6.8), 300 mM KCl, 0.5 mM MgCl_2_, 0.5 mM EGTA, 1 mM NaHCO_3_, and 1 mM DTT. Each sample was treated with an appropriate amount of lambda phosphatase (LP) supplemented by MnCl_2_ and Protein MetalloPhosphatases, as detailed in the manufacturer's protocol (Catalog #P0753L, New England Biolabs, Ipswich, MA). Following treatment, samples were kept at room temperature for 30 minutes and then left at 4°C overnight to allow complete dephosphorylation of proteins. For analyzing the phosphorylation status of proteins, equal volumes (10µl) of various samples were loaded and run on a Mini-PROTEAN Tris-Tricine precast SDS-gels (Bio-Rad, Hercules, CA) following which the gel was treated with Pro-Q diamond stain and destained as described in the manufacturer's manual (P33300 and P33310; Life Technologies, Carlsbad, CA). The phosphorylation status of proteins on the gel was visualized by irradiating it with a fluorescent light following which the gel was stained using Coomassie brilliant blue (R-250; Bio-Rad Laboratories, Hercules, CA) to visualize the total protein amounts. Densitometric analysis of the resulting protein profiles was performed using the ImageJ software (available on the National Institutes of Health website; <https://imagej.nih.gov/ij/> ) to compare the phosphorylation status of proteins among wild-type and R403Q myosin before and after treatment with LP.

**SUPPLEMENTAL RESULTS**

**Effect of Ionic Strength on Single Turnover Kinetics in Myofibrils**

To substantiate our findings in STFs, we also investigated the effect of increasing KCl concentration on the myosin SRX population in a more complicated and physiologically relevant system, namely rabbit cardiac myofibrils (RcMF). Similar to STFs (see Fig. 2 in the main article), the myosin SRX population in RcMF demonstrated a more substantial reduction in response to increasing ionic strength (Fig. S2). For example, increasing KCl concentration from 15 to 100 mM resulted in a net decrease in the SRX population by 26% in RcMF (Fig. S2), while this was 29% in PcSTF (see Fig.2 in the main article). Collectively, these findings suggest that the reconstructed thick filaments can effectively mimic the structural environment similar to the native setting.

#### **Phosphorylation of myosin proteins**

The proteins that undergo phosphorylation in myosin are the regulatory light chain (RLC) and the essential light chain (ELC). A representative gel is shown in Fig. S2 confirms no noticeable differences in the phosphorylation status of RLC and ELC among wild-type and R403Q samples before treatment with LP. To be sure that the results presented in this study are not affected by small differences in the phosphorylation of RLC/ELC between groups, we dephosphorylated samples using LP. Fig. S3 confirms near-complete dephosphorylation in both WT and R403Q myosin.

**Table S1.** Parameters derived using single ATP turnover experiments in various myosin groups following changes in ionic strength (KCl)

| **Parameter** | **KCl concentration** | **PcSTF** | **BcS1** | **BcHMM** |
| --- | --- | --- | --- | --- |
| SRX population,  A_slow_ (%) | 15 mM  30 mM  100 mM  150 mM | 44.6±0.73  26.0±0.77  16.0±3.79  10.1±0.29 | 23.5±2.01  23.3±4.35  23.2±1.75  19.5±1.86 | 30.9±4.78  22.7±0.44  13.4±0.27  11.4±1.06 |
| DRX ATPase rate, *k*_fast_ (s^-1^) | 15 mM  30 mM  100 mM  150 mM | 0.012±0.0001  0.012±0.0001  0.012±0.0003  0.013±0.0007 | 0.013±0.0003  0.013±0.0008  0.009±0.0002  0.008±0.0001 | 0.011±0.0005  0.011±0.0002  0.011±0.0001  0.010±0.0001 |
| SRX ATPase  Rate, *k*_slow_ (s^-1^) | 15 mM  30 mM  100 mM  150 mM | 0.003±0.00005  0.002±0.00015  0.003±0.00062  0.002±0.00008 | 0.0020±0.0002  0.0020±0.0018  0.0012±0.0003  0.001±0.00009 | 0.0018±0.0007  0.0014±0.0002  0.0022±0.0001  0.0020±0.0002 |

Nomenclature is as follows: PcSTF refers to porcine cardiac synthetic thick filaments; BcS1, bovine cardiac myosin sub-fragment S1; BcHMM, bovine cardiac heavy meromyosin; A_slow_, amplitude of myosin SRX population; *k*_fast_ and *k*_slow_ refer to observed ATP turnover rates of myosin population in DRX and SRX states, respectively. Data are reported as mean±SEM (*n*≥4 from two experiments for each group).

**Table S2.** Parameters derived using single ATP turnover experiments in various myosin groups in response to various experimental factors

| **Parameter** | **KCl concentration** | **PcSTF-Phos** | **PcSTF-Dephos** |
| --- | --- | --- | --- |
| SRX population,  A_slow_ (%) | 30 mM | 25.9±0.77 | 45.7±3.7 |
| DRX ATPase rate, *k*_fast_ (s^-1^) | 30 mM | 0.012±0.001 | 0.019±0.002 |
| SRX ATPase  Rate, *k*_slow_ (s^-1^) | 30 mM | 0.002±0.0001 | 0.005±0.0004 |

| **Parameter** | **KCl concentration** | **PcSTF-WT** | **PcSTF-R403Q** |
| --- | --- | --- | --- |
| SRX population,  A_slow_ (%) | 30 mM  100 mM | 45.7±3.7  13.5±2.3 | 25.7±2.5  16.8±3.2 |
| DRX ATPase rate, *k*_fast_ (s^-1^) | 30 mM  100 mM | 0.020±0.002  0.013±0.001 | 0.020±0.001  0.014±0.001 |
| SRX ATPase  Rate, *k*_slow_ (s^-1^) | 30 mM  100 mM | 0.004±0.0002  0.003±0.0007 | 0.003±0.0002  0.002±0.0002 |

| **Parameter** | **ATP vs. ADP** | **PcSTF-WT** | **PcSTF-R403Q** | **BcS1** |
| --- | --- | --- | --- | --- |
| SRX population  A_slow_ (%) | ATP chase  ADP chase | 45.7±3.7  17.8±1.7 | 25.7±2.5  17.7±1.1 | 20.2±3.5  17.2±2.7 |
| DRX ATPase rate  *k*_fast_ (s^-1^) | ATP chase  ADP chase | 0.02±0.002  0.035±0.005 | 0.02±0.001  0.018±0.001 | 0.014±0.001  0.014±0.001 |
| SRX ATPase rate  *k*_slow_ (s^-1^) | ATP chase  ADP chase | 0.005±0.0004  0.003±0.0003 | 0.003±0.0003  0.0015±0.0003 | 0.002±0.0004  0.0013±0.0002 |

Nomenclature is as follows: BcSTF and PcSTF refer to bovine and porcine cardiac synthetic thick filaments, respectively; A_slow_, is the amplitude of myosin SRX population; *k*_fast_ and *k*_slow_ refer to observed ATP turnover rates of myosin population in DRX and SRX states, respectively. Data are reported as mean±SEM (*n*≥4 from two experiments for each group).

**Table S3.** Mavacamten (MAVA) IC_50_ (μM) for different parameters derived using single ATP turnover experiments in various myosin groups in response to various experimental factors

| **System** | **Conditions** | **SRX population**  **(A_slow_)** | **DRX ATPase**  **(*k*_fast_)** |
| --- | --- | --- | --- |
| BcSTF | WT | 0.66±0.05 | 0.31±0.05 |
| BcS1 | WT | 2.16±0.19 | 0.67±0.09 |
| BcHMM | WT | 1.17±0.13 | 0.54±0.08 |

| **System** | **Conditions** | **SRX population**  **(A_slow_)** | **DRX ATPase**  **(*k*_fast_)** |
| --- | --- | --- | --- |
| BcSTF | ATP  ADP | 0.63±0.06  4.35±0.64 | 0.41±0.06 1.14±0.20 |
| BcSTF | α-cardiac  β-cardiac | 1.83±0.25  1.03 ±0.11 | 0.61±0.08  0.35±0.06 |
| PcSTF | WT  R403Q | 1.76±0.22  2.00±0.28 | 0.29±0.02 0.30±0.02 |

Nomenclature is as follows: BcSTF and PcSTF refer to bovine and porcine cardiac synthetic thick filaments, respectively; A_slow_, is the amplitude of myosin SRX population; *k*_fast_ refer to observed ATP turnover rates of myosin population in DRX state; IC_50_ is the concentration of MAVA required to attain the half-maximal change in a given parameter. Data are reported as mean±SEM (*n*≥4 from two experiments for each group).

**SUPPLEMENTAL FIGURES**

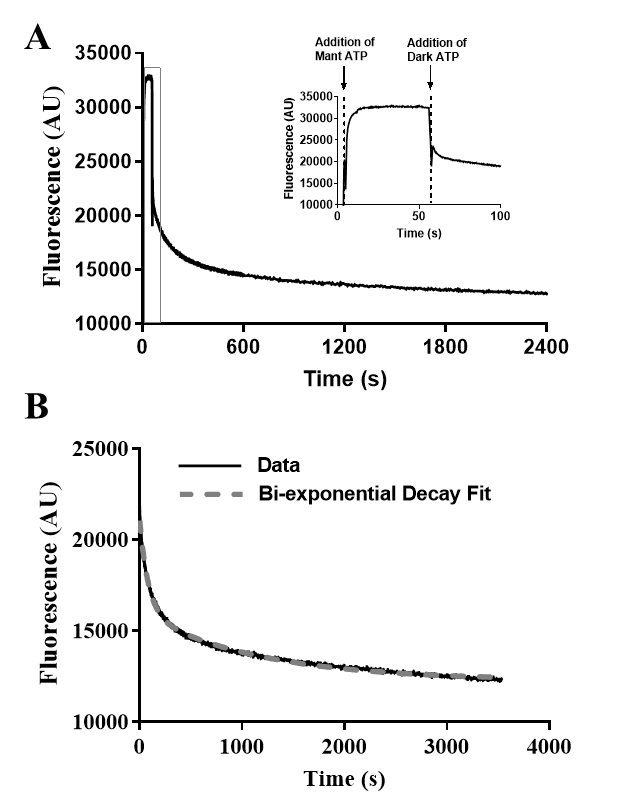

**Figure S1.** Representative trace showing the timing of events in the single mant-ATP turnover experiment. (A) Time course of fluorescence showing mant-ATP binding followed by the unbinding of mant-nucleotides during chase with non-fluorescent ATP. The inset of panel A shows an expanded view of the portion highlighted in the main panel, showing the time points at which mant-ATP and non-fluorescent ATP were added. (B) Full-time course (60 minutes) of decay phase during the chase of mant-nucleotides with non-fluorescent ATP. The decay profile characteristically depicts two phases, a fast phase followed by a slow phase. Each trace was fitted to four different parameters using a bi-exponential decay function to separate into fast and slow phases — A_fast_, *k*_fast_, A_slow_, and *k*_slow_ . A represents the % amplitude and *k* represent the observed ATP turnover rate of each phase. The dotted line in panel B represents the bi-exponential fit to the original trace in black. The values of A_fast_, *k*_fast_, A_slow_, and *k*_slow_ in this representative trace are 72.6%, 0,012 s^-1^, 27.4%, and 0.001 s^-1,^ respectively.

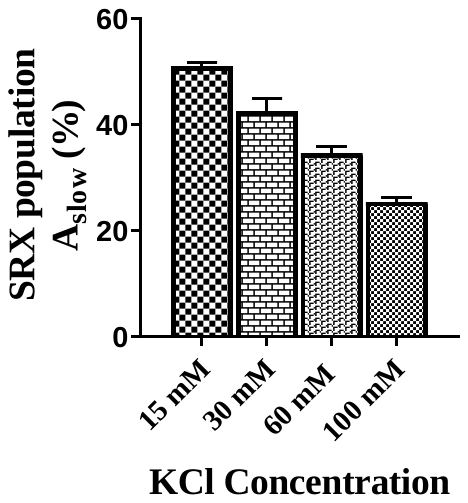

**Figure S2.** Effect of ionic strength (KCl) on the myosin SRX population, A_slow_, in rabbit cardiac myofibril (RcMF) preparations. Changes in myosin SRX population in RcMF observed here are compatible with those in porcine cardiac synthetic thick filaments (PcSTF; see Fig. 2 in the main article). All parameters are reported on the absolute scale. Data are expressed as mean±SEM (*n*=4 for each).

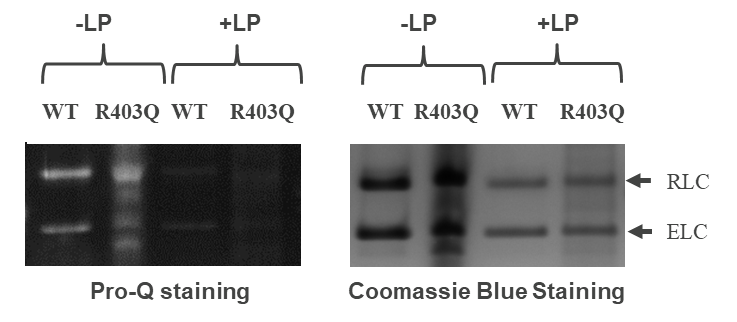

**Figure S3.** SDS-gel analysis of the phosphorylated proteins before and after treatment with lambda phosphatase (LP). Left panel shows Pro-Q stained gel showing the phosphorylation status of proteins in wild-type and R403Q myosin samples before and after treatment with LP, while the right panel shows the total amount of respective proteins following Coomassie blue staining. The phosphorylation levels of RLC and ELC in wild-type and R403Q myosin samples substantially decreased following treatment with LP.

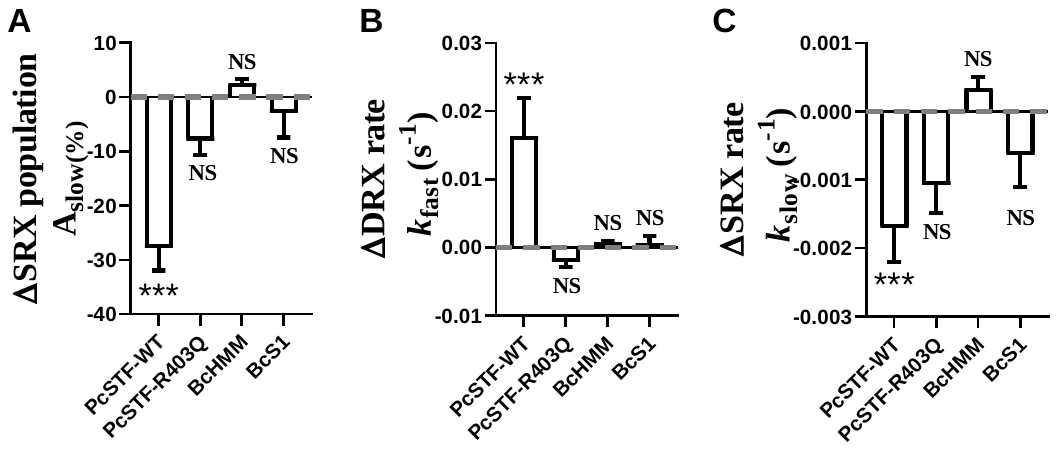

**Figure S4.** Effect of ATP chase versus ADP chase on the parameters derived from single ATP turnover experiments in various myosin systems. Bar graphs showing the effect of ADP chase relative to the ATP chase in PcSTF-WT, PcSTF-R403Q, BcHMM, and BcS1 on (A) A_slow_ (B) *k*_fast_ and (C) *k*_slow_. BcS1 refers to bovine myosin subfragment-1, BcHMM refers to bovine cardiac heavy meromyosin. PcSTF-WT and PcSTF-R403Q refer to porcine cardiac synthetic thick filaments made of WT and mutant R403Q myosin. The dotted line in gray in each panel corresponds to the parameter values in various myosin systems following ATP chase and the bars represent the relative changes in parameters in respective systems following ADP chase. Statistical differences presented are based on two-way ANOVA and subsequent post-hoc Tukey's pair-wise comparisons (***P*<0.01; ****P*<0.001; NS, not significant). Data are expressed as mean±SEM (*n*≥6 for each).

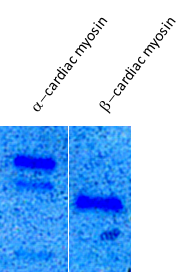

Left

atrial

Left

ventricular

α-cardiac myosin

β-cardiac myosin

**Figure S5.** Native PAGE gel showing the distribution of myosin isoforms in bovine left atrial and bovine left ventricular muscles. As shown, bovine left atrial tissue predominantly expresses an α-cardiac myosin isoform that migrates slower on the gel, while bovine left ventricular tissue predominantly expresses β-myosin isoform that migrates faster on the gel.
